# Females exhibit smaller volumes of brain activation and lower inter-subject variability during motor tasks

**DOI:** 10.1101/2023.04.24.536726

**Authors:** Justin W. Andrushko, Shie Rinat, Eric D. Kirby, Julia Dahlby, Chelsea Ekstrand, Lara A. Boyd

**Affiliations:** Department of Physical Therapy, Faculty of Medicine, University of British Columbia, Vancouver, Canada; Djavad Mowafaghian Centre for Brain Health, University of British Columbia, Vancouver, Canada; Faculty of Individualized Interdisciplinary Studies, Simon Fraser University, Burnaby, BC, Canada; Department of Neuroscience, University of Lethbridge, Lethbridge, Alberta, Canada

**Keywords:** Sex differences, functional magnetic resonance imaging, somatotopic motor mapping

## Abstract

Past work has shown that brain structure and function differ between females and males. Males have larger cortical and sub-cortical volume and surface area (both total and subregional), while females have greater cortical thickness in most brain regions. Functional differences are also reported in the literature, yet to date little work has systematically considered whether patterns of brain activity indexed with functional magnetic resonance imaging (fMRI) differ between females and males. Recently, an attempt to modernize and improve our understanding of the homunculus, a somatotopic representation of the human body within the brain, was made. Still, in the decades since it was introduced, no research has considered sex differences in the somatotopic representation. The current study sought to remediate this issue by employing brain somatotopic motor mapping from a previously published and openly available dataset. We tested differences in patterns of functional brain activity associated with 12 voluntary movement patterns in females versus males. Results suggest that females exhibited smaller volumes of brain activation across all 12 movement tasks, and lower patterns of variability in 10 of the 12 movements. We also observed that females had greater cortical thickness, which is in alignment with previous analyses of structural differences. Overall, these findings provide a basis for considering biological sex in future fMRI research and provide a foundation of understanding differences in how neurological pathologies present in females vs males.

## Introduction

Females and males have anatomical and functional differences in their nervous systems [1–5]. These differences, attributed to differential sex chromosome expression and sex hormones as early as in utero [6–8], manifest in lateralization, motor planning, motor skill, language, memory, and spatial ability [9–14]. However, our understanding of how biological sex affects the somatotopic representations of the body within the brain is limited.

The homunculus, a somatotopic representation of the human body within the sensory and motor cortex, was first introduced in the late 1930s [15]. It maps out the cortical areas responsible for processing information from different body parts, where adjunct body parts are represented by adjacent brain regions. Although there has been a push to modernize our understanding of the classic sensorimotor “homunculus” [16–20], most studies have not considered sex differences. When sex has been included as a factor, the focus was largely on the sexual and reproductive dimorphisms and their respective neuronal representations [20–22].

A comparison of female and male brain structure reveals differences in volume, cortical surface area, cortical thickness, and structural variability [23]. Previous research suggests that males have significantly greater grey matter volume in areas such as the amygdala, temporal pole, fusiform gyrus, putamen, and premotor cortex [3,4]. Further, a large meta-analysis of 16,683 healthy participants found that males exhibit greater subcortical volume as compared to females, with the largest differences found in the thalamus and pallidum, bilaterally [5]. Males also exhibit greater cortical surface area. Importantly, males not only exhibit greater mean differences in subcortical volume and cortical surface area as compared to females, but also significantly greater variability in these measures [5,23]. This variability in surface area is most pronounced in motor related regions such as the pallidum, right inferior parietal cortex, and paracentral region [5]. This suggests that, compared to females, males have less consistent brain structure, particularly in motor related regions.

While most studies suggest greater total and subregional volume in males, some research found regions with greater relative volume in females. When controlling for total brain volume, these studies suggest females show greater grey matter volume than males in prefrontal and superior parietal cortices, as well as the superior temporal sulcus, orbitofrontal cortex, and posterior insula [3,4]. Females also exhibit greater cortical thickness compared to males [5,23]. Further, Wierenga et al. [5] noted regional differences in cortical thickness between females and males; females had greater cortex thickness in 38 of 68 regions, which were primarily in the frontal and parietal cortices. These structural differences suggest that brain function may also differ by sex.

Sex related differences may affect the efficiency of neural activation patterns, with females potentially having more localized and precise, and males having more distributed and varied, neuronal processing. Several sex differences in brain activation have been observed with functional magnetic resonance imaging (fMRI) across a range of tasks and resting-state brain activity [12,24,25]. However, to date, little work has considered whether sex differences affect somatotopic sensorimotor mapping across the entire brain. Researching sex specific patterns of brain activation with respect to area and variability across a range of motor tasks may shed light on key differences in brain function. For instance, in fMRI research, a common approach to investigate brain activation patterns is through region of interest (ROI) analyses. However, using ROIs that are derived from an atlas that is not sex-specific could potentially lead to data selection bias that does not fully capture differences in functional organization in females or males.

Using fMRI whole brain somatotopic motor mapping from a previously published and openly available dataset [[26], https://openneuro.org/datasets/ds004044/versions/2.0.3], the current study in-vestigated the differences in cortical activation patterns between females and males during a variety of motor tasks. Based on previous observations of structural differences, we hypothesized that across movement tasks, females would exhibit lower variability and smaller volumes of brain activation than males, reflecting more focal neural processing. This concerted effort to further quantify the anatomical localization and functional differences will aid our understanding of the inherent differences in somatotopic maps between the sexes, and inform ROI selection strategies for future fMRI research by illustrating whether sex specific somatotopic brain parcellations are needed.

## Methods

### Participants

A total of 68 neurologically intact right-handed young adults (age range 19-29 years) were recruited and screened for the Ma *et al.* [26] study. The original study from which these data were collected was approved by the Institutional Review Board at Beijing Normal University [26], and writen informed consent was provided by the participants.

### Motor tasks

Twelve voluntary movement patterns were each performed for 16 seconds twice during every fMRI scan. A total of six fMRI scans were acquired for each participant, resulting in 12 event blocks for each of the 12 movement types (i.e.,192 seconds of task evoked data for each movement). These movement conditions included bilateral toes, ankles, fingers, wrists, forearms, upper arms, and eyes, in addition to jaw, lip, and tongue movements, and separate unilateral movements for the left and right legs. Detailed movement instructions can be found in table 1. All movements were performed at a self-selected pace between 0.5 and 5 Hz. Importantly, each participant underwent two behavioural training sessions before undergoing the MRI portion of the study. These sessions involved watching a video tutorial on how to perform each of movements, followed by lying supine and practicing each of the movements under experimenter supervision. During these training sessions feedback was given to ensure mastery of movement patterns, magnitude, and speed, while avoiding head or unrelated body movements. To proceed to the MRI portion of the study, movement performance was evaluated and verified by two experimenters. For further details about the nature of the experiment and motor tasks please review the published dataset manuscript [26].

**Table 1.** Movement conditions and instructions

| **Body movements** | **Movement patterns and instructions** |
| --- | --- |
| **Eyes** | Blink or saccade eyes. |
| **Jaw** | Bite or twist jaws. |
| **Lips** | Expand and contract the lips with the teeth being bitten and tongue still. |
| **Tongue** | Circular tongue with the teeth being bitten and lips closed. |
| **Upper arms** | Lift and lower the upper arms (maximum 20°) with the upper arms, forearms, wrist, and fingers straight. |
| **Forearms** | Flex and release both forearms at the elbow (maximum 20°) with the wrists and fingers straight. |
| **Wrists** | Pitch and roll both wrists with clenched fists. |
| **Fingers** | Clench and loose both fists. |
| **Left leg** | Lift and lower the left leg (maximum 10°) with the leg, ankle, and toes straight. |
| **Right leg** | Lift and lower the right leg (maximum 10°) with the leg, ankle, and toes straight. |
| **Ankles** | Dorsiflex and release both ankles. |
| **Toes** | Flex and extend the toes from both feet. |
| **Rest** | Fixate on the dot presented in the center of the screen. |

### Preprocessing structural MRI data

The T1-weighted (T1w) anatomical scans were preprocessed using FSL’s [27] anatomical preprocessing script *fsl_anat*. Briefly, this script performs image reorientation to align with standard space template coordinates, crops the T1w image in the z-direction to remove the neck and other non-brain aspects of the image, implements bias-field correction using FSL’s FAST [28], and then extracts the brain using a non-linear reverse transformation of a standard-space brain mask.

### First level functional MRI processing

The previously preprocessed and manually denoised fMRI data (for more information regarding the original preprocessing and denoising please review Ma et al. [26]) were linearly (ridged body + affine) registered to the bias-field corrected and brain extracted T1w image and then non-linearly (rigid + affine + deformable syn) registered to the MNI152_2mm brain template using antsRegistrationSyN and antsApplyTransforms from Advanced Normalization Tools (ANTs) [29,30]. Each of the 12 motor tasks were modelled using the event timing files provided in the downloaded data with a double-gamma function and with a temporal derivative applied to the model in FSL’s FEAT [31]. White matter BOLD signal was not treated as a nuisance regressor in the present work as this remains a controversial technique [32,33,42–48,34–41]. Therefore, to prevent a bias towards grey matter BOLD signal and to truly investigate whole-brain fMRI signal we retained white matter BOLD in our analyses [46].

### Second level fixed effects functional MRI processing

Since each participant performed the same tasks over six separate runs, a fixed effects analysis was carried out in FEAT [31] prior to performing group-level analyses. The fixed effects analysis was used to average the parameter estimates of the task-based activations across the six runs to create a single activation map and effect size for each motor task for every participant. These averaged parameter estimates were then carried forward for group level analyses in FEAT [49].

### Structural MRI analysis

T1w images for each participant were segmented using Fastsurfer [50]. Overall brain volume without ventricles was first analyzed to determine if there were differences between females and males in brain size. Next, cortical surface area, cortical thickness, and grey matter volume for each hemisphere were extracted for analysis and normalized as a ratio to overall brain volume on a participant-by-participant basis to account for differences in brain size. We then compared each of these normalized metrics for each hemisphere between females and males separately using two-tailed independent samples *t*-tests implemented using the *permutation_test* module from SciPy stats in Python 3.10 [51] with 20,000 permutations. A manual Bonferroni correction to adjust for multiple comparisons was performed and a result was deemed to be significant if the *p*-value was ≤ 0.007 (α = 0.05/7).

### Group level functional MRI analysis

For group-level analyses, two one-sample *t*-tests were performed to compute spatial maps for males and females separately. Additionally, two one-tailed independent samples *t*-tests were carried out to determine if there were differences in the somatotopic representations of each of the 12 motor movements between male and females (*t*-test #1: females > males; *t*-test #2: males > females contrasts). Non-parametric permutation testing was performed on each contrast using *randomise* with 5,000 permutations, and threshold free cluster enhancement was used for statistical inference with a family wise error rate set to α ≤ 0.05 for each contrast.

### Sex difference analyses

To address the question of sex differences for somatotopic motor mapping, two approaches were taken. First, the two *t*-tests previously described (*t*-test #1: females > males; *t*-test #2: males > females contrasts) were assessed and reported when a significant contrast was detected (figure 1). Second, to further investigate sex differences, the mean activation map for each movement was computed separately for males and females and the spatial maps were correlated to assess their degree of similarity. To do this, for males and females the group-level *p*-value threshold statistical maps (α ≤ 0.05) for each movement condition were temporally merged together using *fslmerge*. Then correlation analyses between the spatial maps was carried out with *fslcc*. The correlation coefficients for same movement conditions between females and males were then reported and the inter-task correlations were disgarded.

**Figure 1.**
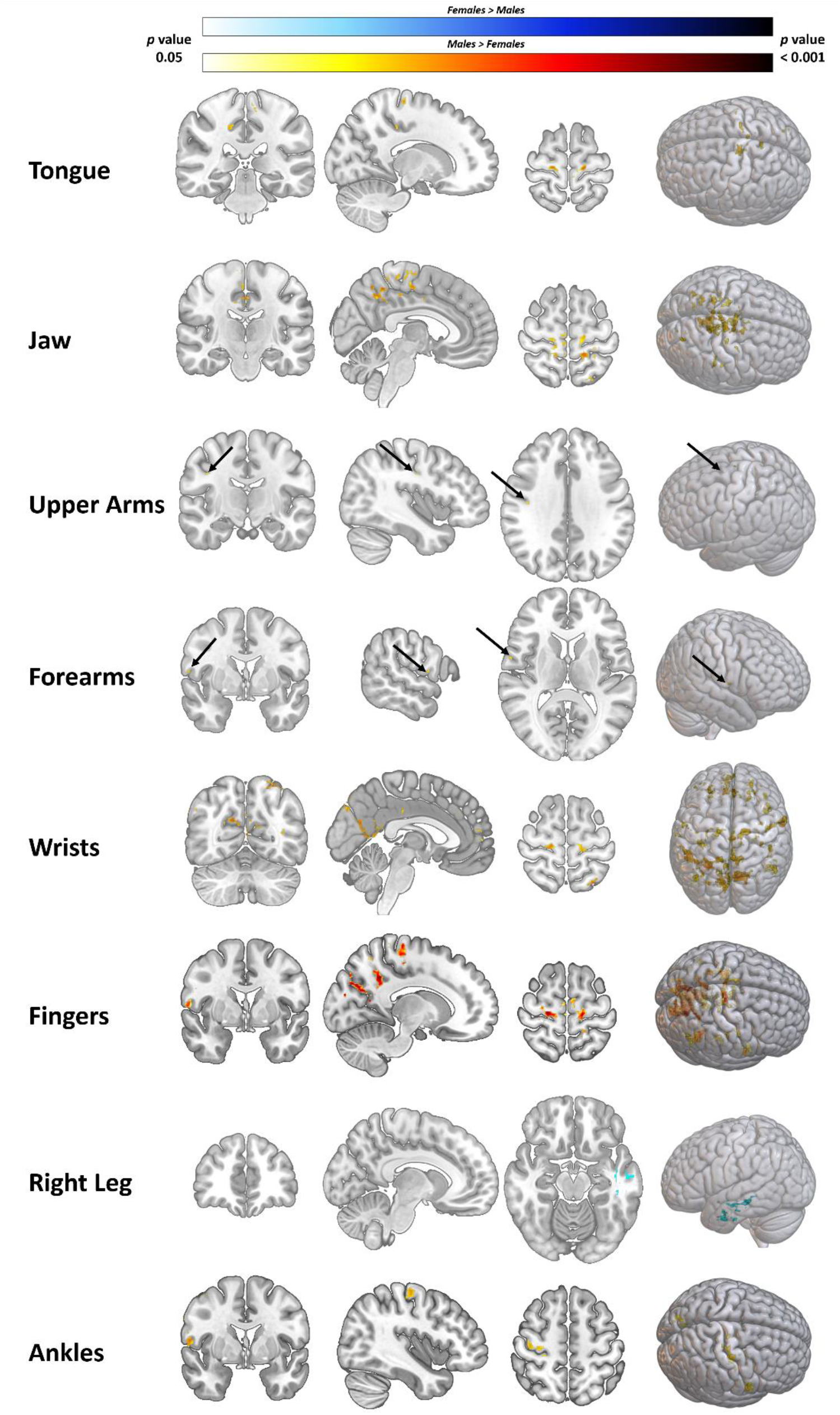
*Sex difference contrasts for 8 significant conditions. Significantly greater BOLD signal for males (Red/Yellow) and females (White/Blue) are plotted. Images are family-wise error corrected p-value maps with a p value threshold of 0.05. Figure was made in MRIcoGL* [52].

### Sex differences in brain activation variability

Variabililty in brain activation was assessed in two ways. First, within group between-subject variability in brain activation was assessed for each motor task. To do this we performed pairwise correlations for both females and males using the raw *t*-statistic image for each participant and calculated the voxel-to-voxel correlation of these images for each pair of participants separately for each group using in-house python code. This process was repeated for each motor task. We then compared the pairwise correlations between females and males for each contrast using two-tailed independent samples *t*-tests implemented using the *permutation_test* module from SciPy stats in Python 3.10 [51] with 20,000 permutations. A manual Bonferroni correction to adjust for multiple comparisons was performed and a result was deemed to be significant if the *p*-value was ≤ 0.004 (α = 0.05/12).

Second, we analyzed brain activation variability within-subject between-runs (6 runs). To do this, we once again perfomed parwise correlations for each participant using the raw t-statistic image for each of the runs. We then calculated the mean correlation coefficient from these pairwise correlations (mean of 15 comparisons). This process was repeated for each of the 12 motor tasks and the rest condition. We then compared the pairwise correlations between females and males for each contrast using two-tailed independent samples *t*-tests implemented using the *permutation_test* module from SciPy stats in Python 3.10 [51] with 20,000 permutations. A manual Bonferroni correction to adjust for multiple comparisons was performed and a result was deemed to be significant if the *p*-value was ≤ 0.004 (α = 0.05/12).

## Results

### Participants

Of the 68 participants from the Ma et al. [26] study, 61 participants were included in the present studies analyses, of which 33 were female (22.58 ± 2.19 years), and 28 were male (23.00 ± 2.34 years). Six participants were excluded from the study because they failed to master the movement patterns as determined by two experimenters prior to the MRI session, and one particpant was exluded from the analyses in this work because they were missing demographics information and their sex could not be determined. Detailed demographic information is provided in original paper [26].

### Sex differences in brain structure

Permutation *t*-testing with 20,000 permutations revealed sex differences in total brain volume [Females: 1,099,719.242 ± 55040.247 mm^3^; Males: 1,230,966.679 ± 110273.860 mm^3^; *t*(59) = -5.912, *p* = 1.00E-04, *d* = -1.480]. There was also a significant difference in normalized cortical thickness in the left [Females: 2.18E-06 ± 1.17E-07; Males: 1.99E-06 ± 1.75E-07; *t*(59) = 4.950, *p* = 1.00E-04, *d* = 1.251] and right [Females: 2.19E-06 ± 1.17E-07; Males: 1.99E-06 ± 1.70E-07; *t*(59) = 5.268, *p =* 1.00E-04, d = 1.333] hemispheres with females having thicker cortex. However, no differences were observed for normalized grey matter volume in the left [Females: 0.211 ± 0.008; Males: 0.214 ± 0.007; *t*(59) = 1.451, *p* = 0.155, *d* = -0.376], or right [Females: 0.212 ± 0.008; Males: 0.215 ± 0.007; *t*(59) = - 1.740, *p* = 0.089, *d* = -0.451] hemisphere, or normalized surface area in the left [Females: 0.075 ± 0.002; Males: 0.075 ± 0.002; *t*(59) = 0.653, *p* = 0.509, *d* = 0.165], or right [Females: 0.075 ± 0.002; Males: 0.075 ± 0.002; *t*(59) = 0.653, *p* = 0.509, *d* = 0.165] hemisphere. These findings suggest that although males have larger overall brains, females exhibited thicker cortex in both hemispheres.

### Sex differences for group-level contrasts

Significant differences were found in males > females contrasts in seven out of the 12 movements including tongue, jaw, upper arms, forearms, wrists, fingers, and ankles (figure 1; supplementary data tables S1-S7). The only contrast that showed a significant females > males difference was right leg movements (data table S8).

### Sex differences for group-level spatial maps

Across all 12 movement tasks, males (mean across 12 tasks: 36,632.58 ± 14,469.06 voxels) exhibited larger volumes of activation compared to females (mean across 12 tasks: 21,315.33 ± 11,738.90 voxels), which equates to males having a mean 92.68 ± 66.81% larger volumes of activation across the whole brain (figure 2; table 8). The spatial maps between females and males were moderately correlated [53] with a mean Pearsons correlation strength *r* = 0.61 ± 0.06, and a range across the 12 tasks from *r* = 0.48 for upper arms movements to *r* = 0.70 for left leg movements (table 2).

**Figure 2.**
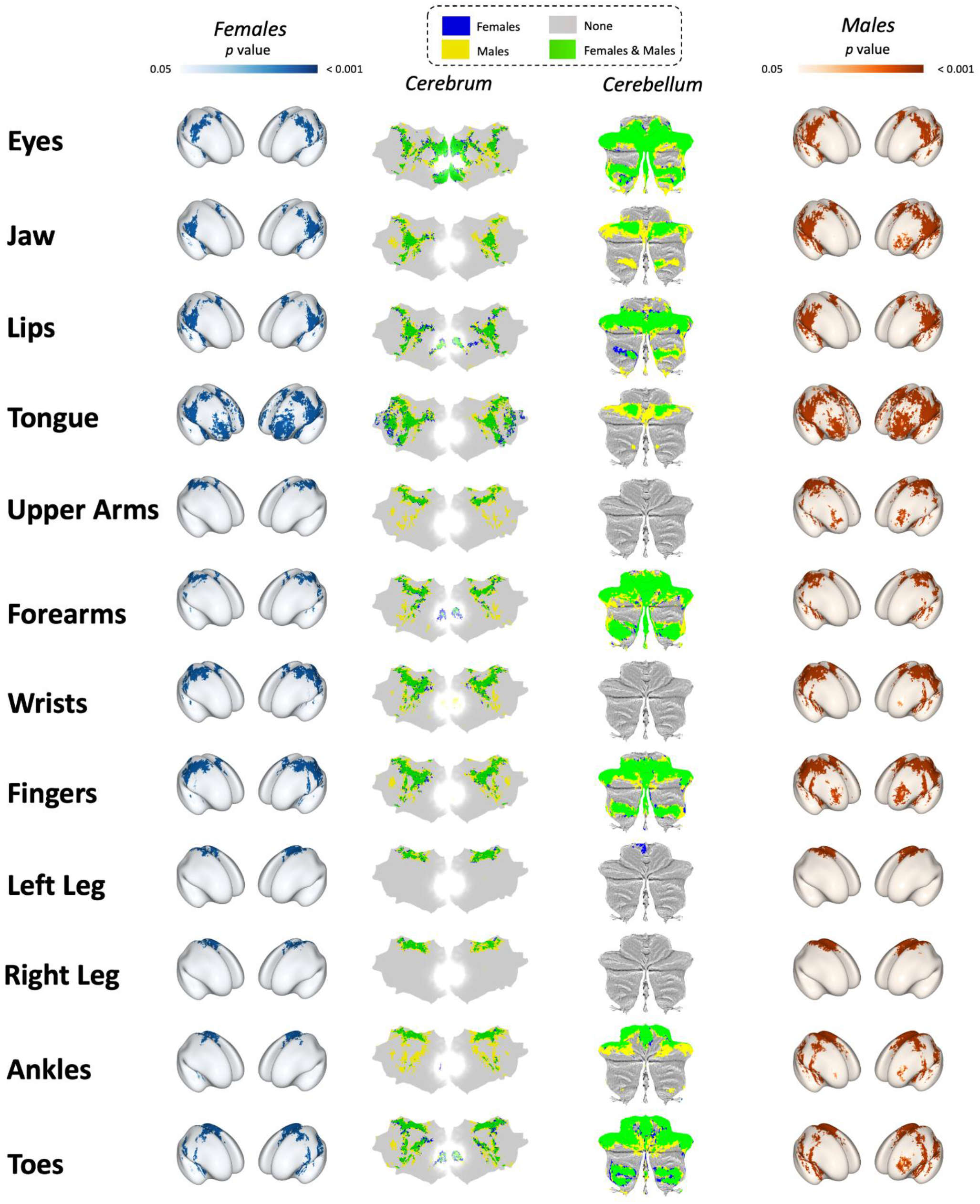
*P-value spatial maps for all 12 movement tasks. Female spatial maps are in blue on the left, male spatial maps are in red/orange on the right. Cerebral and cerebellar flatmaps showing unique and overlap are in the centre two columns respectively. Cerebral inflated brains and flatmaps were made in pycortex* [54]*, and cerebellar flatmaps were made in SUIT* [55].

**Table 2.**
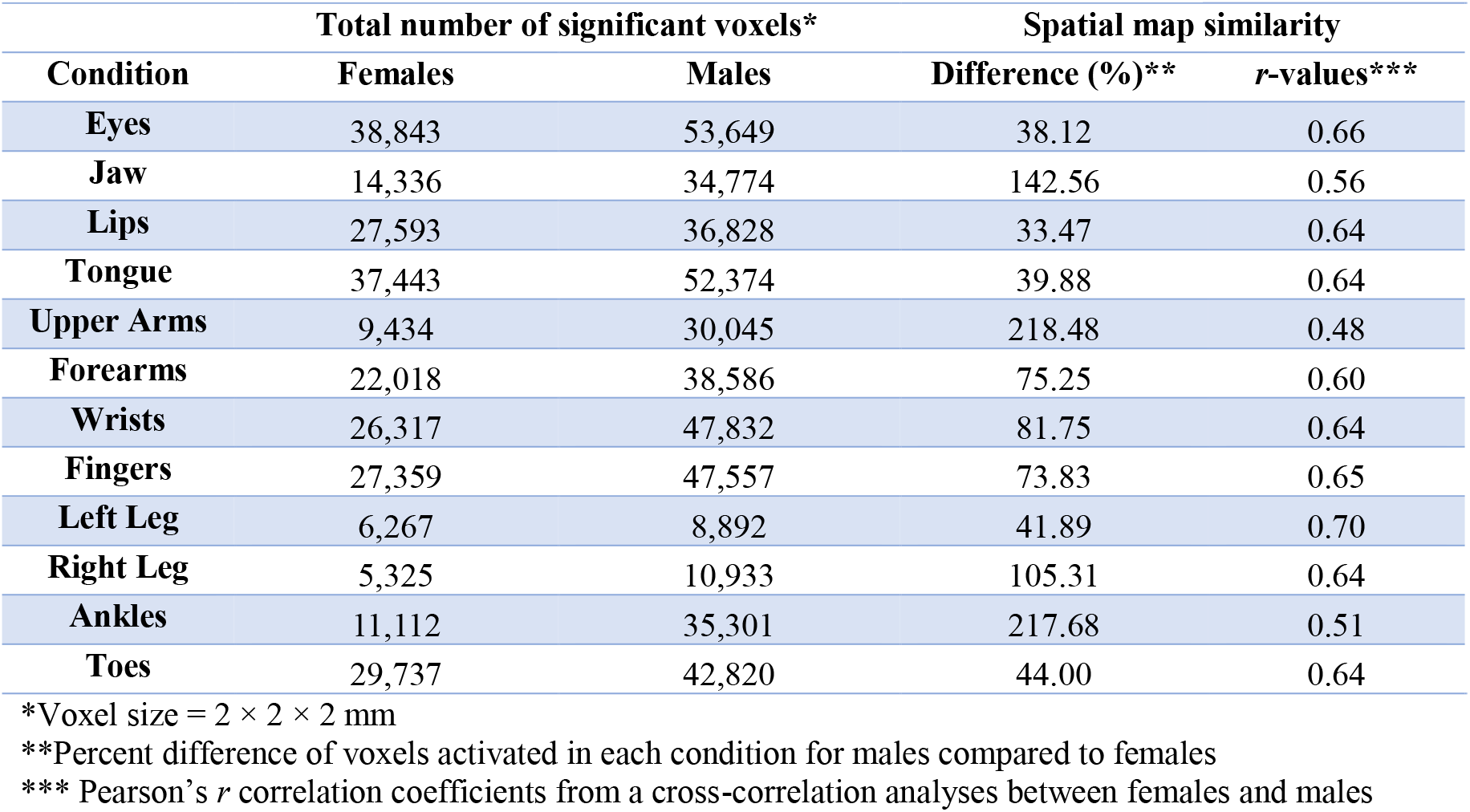
Group-level whole brain spatial maps and correlations between sexes

| Condition | Total number of significant voxels* |  | Spatial map similarity |  |
| --- | --- | --- | --- | --- |
|  | Females | Males | Difference (%)** | r-values*** |
| Eyes | 38,843 | 53,649 | 38.12 | 0.66 |
| Jaw | 14,336 | 34,774 | 142.56 | 0.56 |
| Lips | 27,593 | 36,828 | 33.47 | 0.64 |
| Tongue | 37,443 | 52,374 | 39.88 | 0.64 |
| Upper Arms | 9,434 | 30,045 | 218.48 | 0.48 |
| Forearms | 22,018 | 38,586 | 75.25 | 0.60 |
| Wrists | 26,317 | 47,832 | 81.75 | 0.64 |
| Fingers | 27,359 | 47,557 | 73.83 | 0.65 |
| Left Leg | 6,267 | 8,892 | 41.89 | 0.70 |
| Right Leg | 5,325 | 10,933 | 105.31 | 0.64 |
| Ankles | 11,112 | 35,301 | 217.68 | 0.51 |
| Toes | 29,737 | 42,820 | 44.00 | 0.64 |

\*Voxel size = $2 \times 2 \times 2$ mm
\*\*Percent difference of voxels activated in each condition for males compared to females
\*\*\* Pearson's $r$ correlation coefficients from a cross-correlation analyses between females and males

### Sex differences in brain activation variability

Using permutation testing to assess sex differences in correlations derived from whole-brain voxelwise *t*-statistics for each contrast, we found that females had significantly higher between subject consistency in 10 of the 12 motor tasks (table 3; *p* < 0.004, two-tailed, 20,000 permutations). However an analysis of within subject between run variability resulted in no differences between sex for any of the motor tasks (table 4; two-tailed, 20,000 permutations), suggesting that there were no differences between females and males in how consistent their patterns of brain activation were across the six runs for each of the motor tasks.

**Table 3.** Permutation testing of within group, between subject whole-brain voxelwise brain activation correlations

| Mean Correlation ± Standard Deviation | | | |
| --- | --- | --- | --- |
| Condition | Females | Males | Independent samples t-tests |
| Eyes | 0.259 ± 0.085 | 0.226 ± 0.064 | $t(904) = 6.247, p = 1.00E-04, d = 0.430$ |
| Jaw | 0.228 ± 0.056 | 0.187 ± 0.055 | $t(904) = 11.021, p = 1.00E-04, d = 0.744$ |
| Lips | 0.231 ± 0.066 | 0.201 ± 0.063 | $t(904) = 6.883, p = 1.00E-04, d = 0.465$ |
| Tongue | 0.278 ± 0.069 | 0.264 ± 0.056 | $t(904) = 3.274, p = 0.001, d = 0.224$ |
| Upper Arms | 0.216 ± 0.079 | 0.200 ± 0.061 | $t(904) = 3.483, p = 5.00E-04, d = 0.240$ |
| Forearms | 0.239 ± 0.072 | 0.209 ± 0.047 | $t(904) = 7.253, p = 1.00E-04, d = 0.505$ |
| Wrists | 0.302 ± 0.063 | 0.266 ± 0.062 | $t(904) = 8.530, p = 1.00E-04, d = 0.576$ |
| Fingers | 0.303 ± 0.068 | 0.280 ± 0.066 | $t(904) = 5.179, p = 1.00E-04, d = 0.350$ |
| Left Leg | 0.217 ± 0.085 | 0.183 ± 0.083 | $t(904) = 5.886, p = 1.00E-04, d = 0.398$ |
| Right Leg | 0.199 ± 0.096 | 0.209 ± 0.083 | $t(904) = -1.598, p = 0.108, d = -0.109$ |
| Ankles | 0.216 ± 0.072 | 0.188 ± 0.066 | $t(904) = 5.906, p = 1.00E-04, d = 0.401$ |
| Toes | 0.244 ± 0.070 | 0.184 ± 0.055 | $t(904) = 9.023, p = 1.00E-04, d = 0.620$ |

Note: \* significant at $p < 0.05$ Bonferroni corrected [ $\alpha = 0.004; (0.05/12)$ ], two-tailed, 20,000 permutations.

**Table 4.**
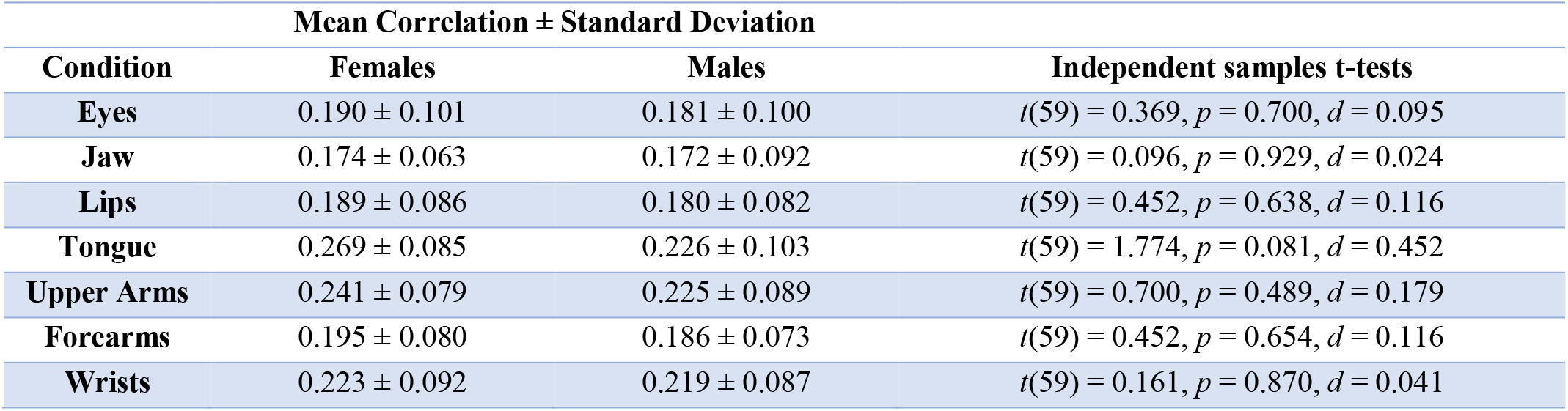
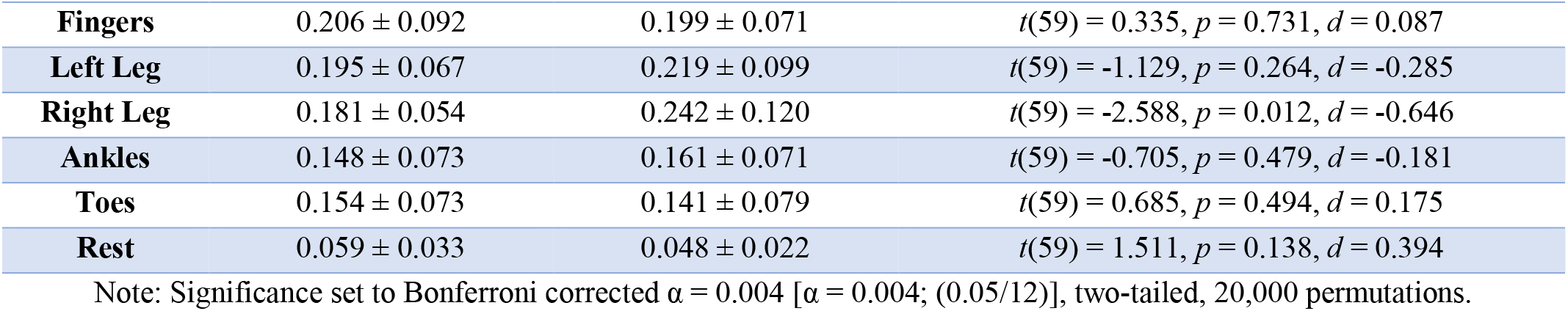
Permutation testing of within subject, between runs whole-brain voxelwise brain activation correlations

| Mean Correlation ± Standard Deviation | | | |
| --- | --- | --- | --- |
| Condition | Females | Males | Independent samples t-tests |
| Eyes | 0.190 ± 0.101 | 0.181 ± 0.100 | $t(59) = 0.369, p = 0.700, d = 0.095$ |
| Jaw | 0.174 ± 0.063 | 0.172 ± 0.092 | $t(59) = 0.096, p = 0.929, d = 0.024$ |
| Lips | 0.189 ± 0.086 | 0.180 ± 0.082 | $t(59) = 0.452, p = 0.638, d = 0.116$ |
| Tongue | 0.269 ± 0.085 | 0.226 ± 0.103 | $t(59) = 1.774, p = 0.081, d = 0.452$ |
| Upper Arms | 0.241 ± 0.079 | 0.225 ± 0.089 | $t(59) = 0.700, p = 0.489, d = 0.179$ |
| Forearms | 0.195 ± 0.080 | 0.186 ± 0.073 | $t(59) = 0.452, p = 0.654, d = 0.116$ |
| Wrists | 0.223 ± 0.092 | 0.219 ± 0.087 | $t(59) = 0.161, p = 0.870, d = 0.041$ |
| **Fingers** | 0.206 ± 0.092 | 0.199 ± 0.071 | $t(59) = 0.335, p = 0.731, d = 0.087$ |
| **Left Leg** | 0.195 ± 0.067 | 0.219 ± 0.099 | $t(59) = -1.129, p = 0.264, d = -0.285$ |
| **Right Leg** | 0.181 ± 0.054 | 0.242 ± 0.120 | $t(59) = -2.588, p = 0.012, d = -0.646$ |
| **Ankles** | 0.148 ± 0.073 | 0.161 ± 0.071 | $t(59) = -0.705, p = 0.479, d = -0.181$ |
| **Toes** | 0.154 ± 0.073 | 0.141 ± 0.079 | $t(59) = 0.685, p = 0.494, d = 0.175$ |
| **Rest** | 0.059 ± 0.033 | 0.048 ± 0.022 | $t(59) = 1.511, p = 0.138, d = 0.394$ |

Note: Significance set to Bonferroni corrected $\alpha = 0.004$ [ $\alpha = 0.004; (0.05/12)$ ], two-tailed, 20,000 permutations.

## Discussion

In this study, we sought to determine if there were sex differences in somatotopic motor maps with a range of movements across the human body. Based on previous research, there are distinct structural differences in cortical organization between females and males, with females having greater cortical thickness [5,23], and males having larger cortical surface area and cortical volume; this is particularly the case for the motor and somatosensory regions [5,23] which are known to have somatotopic organization [56]. Given these previously observed differences, we hypothesized that females would exhibit more focal (i.e., smaller volumes of activation) and consistent (i.e., lower variability between females for each task) patterns of brain activation, compared to males across the 12 different body movements. To answer these questions, we used a freely available fMRI dataset that included a total of 61 participants, of which 33 were females and 28 were males.

### Males generally exhibit larger magnitudes of brain activation

In seven of the movements under investigation (jaw, tongue, upper arms, forearms, wrists, fingers, and ankles) we observed greater BOLD signal in males than females. In contrast, females only exhibited greater activation than males during right leg movements, and no differences were seen with the remaining four movements (eyes, lips, left leg, and toes).

### Females exhibit smaller volumes of brain activation across all movement conditions

Smaller activation maps were observed in females across all 12 motor tasks (on average males had 92.68 ± 66.81% larger activation maps than females, ranging from 33.47% larger for lip movements to 218.48% larger for upper arms movements; table 2, figure 2). These data support our hypothesis that females would have more focal volumes of brain activation given the previous findings that males have larger overall cortical and subcortical volume and surface area [5,23], and provide support for females having better cortical efficiency [57] during simple movements across the body.

### Lower whole-brain activation variability in females during all movement conditions

Based on our novel whole-brain voxelwise correlation analysis, females consistently showed higher correlations between subjects compared to males (table 2); these differences were statistically significant. These findings should be viewed as complementary to the volume of activation data reported in this work, given the smaller volumes of brain activation paired with the lower between subject variability seen in females may be indicative of greater neural efficiency compared to males. Additionally, to determine if there were sex differences for how consistent patterns of brain activation were during the 12 motor tasks, we assessed intra-subject variability for each motor task across the six separate fMRI runs. Based on these intra-subject variability analyses, there were no sex differences and both females and males exhibited similar levels of variability. These analysis were important to ensure any sex differences observed throughout the study were not a product of intra-subject variability differences between sexes.

Our findings contribute to the growing body of evidence indicating that extensive sex differences in brain organization and function exist [3,23,58,59]. Importantly, Ritchie *et al.*, [23] noted that the observed sex differences in cortical measures in their work may be related compensatory interactions between sex-specific hormones and the neural underpinnings that govern human voluntary movement [60–62]. It is plausible that these hormonal differences may impact both behaviour and the structure and function of the human brain, which could explain at least in part the differences observed in the present study.

It is well known that the primary motor cortex is organized somatotopically, with adjacent body parts represented by neighbouring regions of the cortex [56]. Recently, the accuracy of the motor homunculus has been brought into question, with a proposed update to the cortical representation maps [16]. However, little is known about the extent to which this organization may vary between females and males. There is accumulating evidence from recent years suggesting sex differences in brain structure, and specifically in motor areas, including the premotor cortex, putamen, and cerebellum [GM volume higher in males > females [4,23,63]]. Yet, the majority of these findings refer to structural differences in volume, surface area or cortical thickness, with limited data regarding functional imaging differences or their associations with motor behaviour. Greater spatial distributions and/or magnitudes of BOLD signal have been observed, without differences in task performance, for males during verbal and spatial tasks [64], in addition to cognitive and motor tasks [65]. These findings paired with the present work suggest that females often show lower magnitudes, and smaller volumes of brain activation across cognitive-motor tasks.

### Implications of sex differences in functional brain imaging research analyses

Importantly, there are many methods to analyze functional brain imaging data. These analysis and interpretation approaches often rely on previous findings, and as a result neglect to acknowledge sex differences. For example, a ROI based analysis that is derived from an atlas, could prevent true ROI identification if it does not consider sex differences in functional organization of the brain, and is instead a product of group averaging. Some brain atlases that are commonly used in fMRI research for ROI selection include the Harvard-Oxford cortical and subcortical atlases (16 females, 21 males, between the ages of 18-50), the Brainnectome atlas (10 females, 10 males, all right handed, between the ages of 19-25), and the Shaefer atlas (665 males, 905 females, between the ages of 18-35). Each of these atlases were created by mixing female and male participants, which may lead to suboptimal parcellations given our findings that males tend to exhibit larger volumes of BOLD signal across a range of motor tasks. Our data suggests that future studies should consider selection of sex specific ROIs based on functional activation [66–68], when possible.

### Implications of sex differences in neurological impairments

As discussed, a knowledge gap exists regarding the dimorphisms in somatotopic maps between females in males. While filling this gap is generally useful for research purposes, it cannot be understated how important these findings may be for the understanding of the etiology, prognosis, and recovery from injury or disease in females versus males. The incidence and presentation of many neurological conditions and movement disorders differ between sexes, suggesting underlying differences in brain function or structure [69]. For example, females experience increased severity, more impairment, and longer recovery from stroke [70–73]. Although many explanatory factors have been identified, incorporating somatotopic differences into this phenomenon should be discussed. As identified in this research, females use less of their cortex to generate voluntary movements, and do so with significantly lower brain activation variability. Although this may be beneficial in healthy, neurologically intact populations, this approach may be detrimental in the presence of a neurological injury. For example, with stroke, injury to the central nervous system interrupts the normal motor output pathways and leads to motor impairment. Our data showing that females have smaller volumes of cortical activation than males, and lower variability, shows that comparisons against standard ROI atlases may lead to misinterpretation of data and fundamentally limit our understanding of brain function.

## Limitations and future directions

As this study used a publicly available data, our analysis was restricted to the previously collected data. Therefore, we were unable to include gender in our analysis. Recent studies indicate that gender differences in both brain structure and brain function exist [74], and future studies should collect the appropriate data to allow for both sex- and gender-based analyses. Additionally, force output can influence brain activation [75–77], and in the present data there are no behavioural metrics to quantify the movement force, velocity, or quality. This is a noted limitation that future work should consider measuring to better link movement kinematics to cortical measures of activation. However, past work [75] suggests that small differences in force (e.g., between 25% and 50%, or 50% and 75% MVC) are not associated with significant differences in brain activation. Only large differences in movement characteristics (e.g., 25% and 75% MVC) result in activation changes. Such differences are unlikely to occur in this study, as participants were instructed to perform the movements at a comfortable self-selected pace, without any additional resistance. Further, even if individual differences exist, group differences in movement characteristics are not assumed, especially in light of the lack of sex differences in intra-subject variability. In light of our sample size, we presume that such differences, if they exist, will be balanced between the two groups.

## Conclusions

In this study we found significant sex differences in the magnitude of brain activation for eight out of the 12 movement conditions, with males having significantly greater brain activation in seven out of the 12 conditions and females showing greater brain activation in one condition. Interestingly, we observed smaller volumes of brain activation in all 12 movement conditions and significantly lower whole-brain activation variability in 10 of the 12 movement conditions for females. With a cross-correlation analysis to compare the spatial map similarity between sexes, we found that spatial maps were moderately correlated for each movement conditions between sexes. The difference in volume is an important consideration when creating ROIs based on functional localizers, as clear and visible differences can be observed between sexes, and has potentially significant implications for understanding the neurobiology of various neurological impairments in females and males (figure 2). We propose these data provide important insight into the need for sex specific ROIs for functional localization to accurately study brain-behaviour relationships using fMRI, and likely have important clinical implications for understanding impairment severity and rehabilitation strategies for each sex independently.

## Acknowledgements

The authors would like to acknowledge the original authors of the open dataset Sai Ma, Taicheng Huang, Yukun Qu, Xiayu Chen, Yajie Zhang and Zonglei Zhen. Without their support of open science practices, this work would not have been made possible.

## Author contributions

J.W.A. processed and analyzed the data, and assisted in writing the manuscript. S.R. processed and analyzed the data and assisted in writing the manuscript. E.K. created the figures and assisted in writing the manuscript. J.D. assisted with writing the manuscript and approving the final version. C.E. assisted in data analysis and writing the manuscript, and C.E. and L.B. were involved with the overall conception and direction of this study, and edited and approved the manuscript.

