## supplementary data tables for "Females exhibit smaller volumes of brain activation and lower inter-subject variability during motor tasks"

Supplementary Tables

**Table S1.** Jaws, Males > Females

| | | Max Intensity | | | Center of Gravity | | |
|---|---|---|---|---|---|---|---|
| **Voxels** | **Z-Max** | **X** | **Y** | **Z** | **X** | **Y** | **Z** |
| 2070 | 4.89 | 10 | -6 | 38 | 0.361 | -34.3 | 53.7 |
| 143 | 3.93 | -12 | -22 | 78 | -14.9 | -24.4 | 66.7 |
| 80 | 5.03 | 34 | -32 | 22 | 43.3 | -34.3 | 18.2 |
| 33 | 3.98 | 30 | -20 | 62 | 29.3 | -21 | 58.8 |
| 33 | 3.97 | 30 | -10 | 42 | 33.5 | -9.3 | 46.1 |
| 28 | 4.97 | 26 | -74 | 42 | 27.6 | -73.1 | 41.1 |
| 22 | 3.91 | 44 | -26 | 14 | 46.2 | -25.2 | 14 |
| 18 | 4.58 | 20 | -80 | 46 | 19.8 | -79.1 | 49.1 |
| 17 | 3.37 | 10 | -52 | 72 | 11.8 | -52.3 | 73.4 |
| 17 | 3.83 | 16 | -70 | 30 | 14.5 | -71.1 | 32.7 |
| 15 | 3.83 | -12 | -12 | 62 | -9.7 | -10.8 | 63.7 |
| 9 | 3.96 | 36 | -76 | 44 | 35 | -77.3 | 43.8 |
| 8 | 3.59 | -8 | -18 | 62 | -7.26 | -19.2 | 61.8 |
| 6 | 3.82 | 12 | -78 | 46 | 13.3 | -77.4 | 47 |
| 6 | 4.53 | -16 | -82 | 44 | -15.4 | -80.7 | 43.4 |
| 4 | 3.71 | 46 | -16 | 46 | 46.9 | -16.5 | 45.5 |
| 4 | 3.68 | 6 | -72 | 40 | 5.48 | -72.6 | 39.5 |
| 4 | 3.46 | 46 | -64 | 48 | 46 | -65.5 | 47.5 |
| 4 | 3.15 | 38 | -18 | 58 | 38.5 | -18 | 56.5 |
| 3 | 3.78 | 10 | -64 | 32 | 10 | -63.3 | 32 |
| 3 | 3.09 | -18 | -14 | 64 | -18 | -12.7 | 63.4 |
| 2 | 4.09 | 22 | -56 | 20 | 22 | -56 | 19 |
| 2 | 4.05 | 22 | -66 | 38 | 22 | -65 | 37 |
| 2 | 3.77 | -12 | -66 | 28 | -12 | -66 | 29 |
| 1 | 3.41 | 0 | -40 | 38 | 0 | -40 | 38 |
| 1 | 4.04 | 34 | -26 | 42 | 34 | -26 | 42 |
| 1 | 3.8 | 40 | -52 | 38 | 40 | -52 | 38 |
| 1 | 3.87 | 40 | -50 | 48 | 40 | -50 | 48 |
| 1 | 3.33 | 8 | -78 | 50 | 8 | -78 | 50 |
| 1 | 4.37 | 40 | -56 | 18 | 40 | -56 | 18 |
| 1 | 3.01 | 30 | -18 | 50 | 30 | -18 | 50 |
| 1 | 3.72 | 50 | 2 | 52 | 50 | 2 | 52 |
| 1 | 3.79 | 48 | 2 | 56 | 48 | 2 | 56 |
| 1 | 3.9 | 6 | -70 | 26 | 6 | -70 | 26 |
| 1 | 4.97 | 44 | -48 | 62 | 44 | -48 | 62 |
| 1 | 2.54 | 6 | -52 | 68 | 6 | -52 | 68 |

Supplementary Tables

**Table S2.** Tongue, Males > Females

| | | Max Intensity | | | Center of Gravity | | |
|---|---|---|---|---|---|---|---|
| **Voxels** | **Z-Max** | **X** | **Y** | **Z** | **X** | **Y** | **Z** |
| 128 | 4.6 | -16 | -24 | 68 | -13.1 | -25.4 | 66.7 |
| 48 | 3.74 | 2 | -48 | 42 | 4.51 | -48.9 | 44.9 |
| 47 | 4.83 | 18 | -24 | 66 | 15.5 | -24.2 | 68.3 |
| 19 | 3.85 | 8 | -38 | 52 | 11.7 | -38.5 | 57.5 |
| 15 | 4.19 | 18 | -32 | 40 | 15.9 | -32 | 42.1 |
| 15 | 4.46 | 20 | -80 | 46 | 22.8 | -79.6 | 47.9 |
| 14 | 3.92 | 46 | -16 | 44 | 46.3 | -17.1 | 46.2 |
| 9 | 4.82 | 26 | -74 | 42 | 26.5 | -73.1 | 40.8 |
| 4 | 3.6 | -4 | -32 | 40 | -3.01 | -32.4 | 40.5 |
| 3 | 3.94 | -12 | -32 | 50 | -12 | -32.6 | 49.4 |
| 2 | 4.14 | 34 | -82 | 24 | 34 | -81 | 24 |
| 2 | 3.88 | 14 | -90 | 34 | 13 | -90 | 33 |
| 2 | 4.31 | 24 | -66 | 38 | 24 | -66 | 39 |
| 2 | 4.13 | -6 | -28 | 46 | -5.08 | -28 | 46 |
| 2 | 3.31 | -26 | -36 | 62 | -26 | -36 | 61 |
| 2 | 4.51 | 44 | -48 | 62 | 44 | -49 | 62 |
| 1 | 4.77 | 22 | -76 | 16 | 22 | -76 | 16 |
| 1 | 5.24 | 36 | -36 | 20 | 36 | -36 | 20 |
| 1 | 4.19 | 18 | -70 | 30 | 18 | -70 | 30 |
| 1 | 4.64 | 32 | -82 | 34 | 32 | -82 | 34 |
| 1 | 3.88 | 18 | -70 | 34 | 18 | -70 | 34 |
| 1 | 3.8 | 8 | -38 | 36 | 8 | -38 | 36 |
| 1 | 3.86 | -6 | -36 | 46 | -6 | -36 | 46 |
| 1 | 3.32 | 0 | -42 | 50 | 0 | -42 | 50 |
| 1 | 3.77 | -24 | -36 | 54 | -24 | -36 | 54 |
| 1 | 3.32 | 36 | -20 | 54 | 36 | -20 | 54 |
| 1 | 3.68 | 4 | -20 | 56 | 4 | -20 | 56 |
| 1 | 3.79 | 8 | -40 | 58 | 8 | -40 | 58 |

**Table S3.** Upper Arms, Males > Females

| | | Max Intensity | | | Center of Gravity | | |
|---|---|---|---|---|---|---|---|
| **Voxels** | **Z-Max** | **X** | **Y** | **Z** | **X** | **Y** | **Z** |
| 15 | 4.73 | 40 | -12 | 34 | 41.9 | -12.8 | 34 |

| | | | | | | | |
|---|---|---|---|---|---|---|---|
| 5 | 4.56 | 60 | 2 | 10 | 59.2 | 1.62 | 10.8 |
| 5 | 4.53 | 64 | -2 | 32 | 64 | -1.24 | 29.6 |
| 3 | 4.9 | -30 | 26 | 2 | -30.6 | 26.6 | 2 |

**Table S4.** Forearms, Males > Females

| | | Max Intensity | | | Center of Gravity | | |
|---|---|---|---|---|---|---|---|
| **Voxels** | **Z-Max** | **X** | **Y** | **Z** | **X** | **Y** | **Z** |
| 7 | 4.92 | 58 | 0 | 10 | 59.1 | 0.86 | 10.3 |

**Table S5.** Wrists, Males > Females

| | | Max Intensity | | | Center of Gravity | | |
|---|---|---|---|---|---|---|---|
| **Voxels** | **Z-Max** | **X** | **Y** | **Z** | **X** | **Y** | **Z** |
| 1660 | 5.01 | -24 | -84 | 44 | -12.9 | -65.3 | 37 |
| 607 | 5.32 | -10 | 34 | 18 | -4.18 | 44.7 | 13.5 |
| 352 | 4.83 | 46 | -8 | 30 | 52.6 | -1.45 | 20.8 |
| 336 | 4.8 | 4 | -52 | 42 | 4.75 | -40.8 | 38 |
| 229 | 4.75 | -30 | 28 | 56 | -25.3 | 28.4 | 47 |
| 210 | 4.99 | 54 | -60 | 30 | 49.8 | -58.4 | 30.9 |
| 177 | 5.03 | -28 | -6 | -6 | -31.7 | -19.5 | 1.92 |
| 166 | 4.15 | 18 | 46 | 36 | 21.4 | 32.4 | 40 |
| 160 | 4.9 | 8 | -30 | 60 | 14.9 | -26 | 64.2 |
| 94 | 4.19 | 42 | 16 | 32 | 40.5 | 12.5 | 33.6 |
| 93 | 3.92 | -40 | -74 | 28 | -39.6 | -80.6 | 32.1 |
| 74 | 3.85 | -42 | -18 | 32 | -51.6 | -10.8 | 25 |
| 73 | 3.6 | -8 | -24 | 68 | -14.8 | -26 | 64.6 |
| 65 | 4.35 | 28 | -12 | -2 | 29.3 | -4.78 | -6.95 |
| 63 | 3.87 | -38 | -64 | 12 | -39.6 | -68.4 | 16.4 |
| 51 | 3.59 | 46 | -34 | 22 | 46.4 | -30 | 19.7 |
| 33 | 4.15 | -64 | -22 | 34 | -62.7 | -23.5 | 36.5 |
| 27 | 3.69 | -32 | 44 | 12 | -36 | 45 | 16.5 |
| 27 | 3.62 | 38 | 26 | 40 | 37.7 | 27.2 | 40 |
| 26 | 4.81 | 36 | -24 | 4 | 35.7 | -22.6 | 4.71 |
| 24 | 4.59 | 38 | -42 | 18 | 42.1 | -41.9 | 17.6 |
| 21 | 4.78 | -44 | 38 | 32 | -44.5 | 36.1 | 33.6 |
| 19 | 3.88 | 12 | -32 | 48 | 11.9 | -34 | 47.3 |

Supplementary Tables

| | | | | | | | |
|---|---|---|---|---|---|---|---|
| 18 | 4.12 | 2 | 62 | 28 | 2.11 | 61.2 | 32.5 |
| 18 | 3.19 | 32 | 14 | 8 | 35.2 | 12.4 | 8.23 |
| 17 | 4.05 | 32 | 40 | 46 | 31.3 | 38.3 | 47.1 |
| 16 | 3.26 | -16 | 36 | 52 | -12.7 | 36 | 55 |
| 12 | 3 | 28 | -30 | 60 | 26.3 | -31.3 | 59 |
| 9 | 3.07 | 24 | 16 | 46 | 23.6 | 16.8 | 47.5 |
| 8 | 3.8 | 4 | -10 | 6 | 3.5 | -11 | 7.45 |
| 7 | 4.21 | 22 | -82 | 46 | 21.9 | -81.2 | 46 |
| 6 | 3.25 | 48 | -8 | 10 | 49 | -8.35 | 10.6 |
| 6 | 3.65 | -34 | 14 | 46 | -33.7 | 13.3 | 47 |
| 6 | 3.37 | -38 | 4 | 50 | -38 | 5.29 | 50.3 |
| 5 | 3.39 | 68 | -12 | 4 | 68.8 | -13.2 | 3.23 |
| 5 | 4.89 | 44 | -50 | 62 | 41.6 | -50.8 | 63.2 |
| 5 | 3.28 | 2 | 46 | -8 | 2.76 | 47.1 | -6.83 |
| 4 | 3.71 | 34 | 24 | 28 | 33.1 | 23.5 | 28 |
| 4 | 4.08 | 26 | -74 | 42 | 26.4 | -73.5 | 40.5 |
| 4 | 3.63 | 24 | 32 | 54 | 25.4 | 30.6 | 55.5 |
| 4 | 3.22 | -20 | -18 | 54 | -20 | -18.5 | 55.5 |
| 4 | 3.08 | 0 | 68 | 2 | -0.49 | 68 | 1.93 |
| 4 | 2.99 | 2 | 46 | 0 | 1.49 | 47.5 | -0.974 |
| 4 | 3.94 | -66 | -6 | 12 | -66 | -6.49 | 12 |
| 4 | 3.91 | 2 | 20 | 30 | 2 | 20 | 29 |
| 4 | 4.47 | 16 | 24 | 10 | 15.6 | 22.6 | 10.4 |
| 4 | 3.91 | 34 | 18 | 26 | 33.1 | 18 | 25.1 |
| 3 | 3.14 | 62 | -14 | 12 | 62 | -14 | 12 |
| 3 | 3.9 | -64 | -2 | 8 | -63.4 | -0.722 | 7.36 |
| 3 | 2.82 | -8 | 28 | 22 | -8 | 27.3 | 22.7 |
| 3 | 2.49 | 42 | -66 | 20 | 42 | -66 | 20 |
| 3 | 3.93 | -46 | 28 | 42 | -47.3 | 26 | 40.7 |
| 3 | 4.01 | -26 | -10 | 2 | -25.4 | -8.73 | 2 |
| 3 | 3.33 | 12 | -58 | 26 | 11.4 | -58 | 26.6 |
| 3 | 3.67 | -8 | 44 | -10 | -7.34 | 45.3 | -10 |
| 3 | 2.97 | 36 | -52 | 38 | 37.9 | -52 | 38 |
| 3 | 3.15 | 22 | 22 | 62 | 23.3 | 21.4 | 62 |
| 2 | 2.84 | 24 | 16 | 40 | 24 | 17 | 40 |
| 2 | 3.91 | 46 | -22 | 22 | 46 | -21 | 22 |
| 2 | 3.73 | 26 | 42 | -16 | 26.9 | 42 | -16 |
| 2 | 4.36 | -38 | 20 | 54 | -38 | 20.9 | 54 |
| 2 | 3.6 | -4 | 18 | 20 | -4.96 | 18 | 20 |

### Supplementary Tables

| | | | | | | | |
|---|---|---|---|---|---|---|---|
| 2 | 3.29 | -2 | -74 | 56 | -2 | -74 | 55 |
| 2 | 3.37 | -16 | 44 | 50 | -15 | 45 | 50 |
| 2 | 3.08 | 18 | 16 | -10 | 19 | 17 | -10 |
| 2 | 2.98 | -20 | 24 | 48 | -20 | 24 | 49 |
| 2 | 3.19 | -4 | 54 | -10 | -4 | 54 | -9.09 |
| 2 | 3.68 | -18 | -44 | -6 | -17.1 | -44.9 | -5.08 |
| 2 | 3.14 | 56 | -16 | 12 | 56 | -16 | 11 |
| 2 | 2.94 | -54 | -32 | 38 | -54 | -33 | 39 |
| 2 | 3.67 | 56 | -16 | 6 | 56.9 | -16 | 6 |
| 2 | 3.5 | -44 | 6 | 34 | -43 | 6 | 34 |
| 2 | 3.8 | 42 | 24 | 6 | 42 | 24 | 6.97 |
| 1 | 3.62 | 30 | -12 | 8 | 30 | -12 | 8 |
| 1 | 3.38 | -2 | -46 | 36 | -2 | -46 | 36 |
| 1 | 3.35 | -62 | 4 | 8 | -62 | 4 | 8 |
| 1 | 3.24 | 30 | 18 | 8 | 30 | 18 | 8 |
| 1 | 3.25 | -30 | 8 | 38 | -30 | 8 | 38 |
| 1 | 3.14 | -46 | -64 | 40 | -46 | -64 | 40 |
| 1 | 3.34 | -34 | 8 | 40 | -34 | 8 | 40 |
| 1 | 3.43 | -44 | 10 | 40 | -44 | 10 | 40 |
| 1 | 2.97 | -40 | 14 | 40 | -40 | 14 | 40 |
| 1 | 3.54 | 38 | -40 | 24 | 38 | -40 | 24 |
| 1 | 4.37 | 16 | 28 | 4 | 16 | 28 | 4 |
| 1 | 2.87 | 32 | 40 | 42 | 32 | 40 | 42 |
| 1 | 3.15 | -44 | -64 | 44 | -44 | -64 | 44 |
| 1 | 2.64 | 34 | 2 | 4 | 34 | 2 | 4 |
| 1 | 3.62 | 22 | 20 | -6 | 22 | 20 | -6 |
| 1 | 3.87 | -20 | 12 | -6 | -20 | 12 | -6 |
| 1 | 3.42 | 2 | 50 | 44 | 2 | 50 | 44 |
| 1 | 3.97 | -58 | 2 | 10 | -58 | 2 | 10 |
| 1 | 4.18 | -20 | 8 | -8 | -20 | 8 | -8 |
| 1 | 2.84 | 0 | 64 | 10 | 0 | 64 | 10 |
| 1 | 4.07 | -18 | -50 | 28 | -18 | -50 | 28 |
| 1 | 3.49 | 6 | 44 | 14 | 6 | 44 | 14 |
| 1 | 3.64 | -48 | -4 | 16 | -48 | -4 | 16 |
| 1 | 3.9 | -6 | 48 | 50 | -6 | 48 | 50 |
| 1 | 3.93 | 24 | 18 | -12 | 24 | 18 | -12 |
| 1 | 2.8 | -40 | 10 | 52 | -40 | 10 | 52 |
| 1 | 3.97 | -14 | 0 | 20 | -14 | 0 | 20 |
| 1 | 3.26 | 26 | 38 | -14 | 26 | 38 | -14 |

Supplementary Tables

| | | | | | | | |
|---|---|---|---|---|---|---|---|
| 1 | 3.9 | 24 | 10 | -14 | 24 | 10 | -14 |
| 1 | 3.15 | 2 | -32 | 26 | 2 | -32 | 26 |
| 1 | 3.04 | 30 | 30 | 54 | 30 | 30 | 54 |
| 1 | 3.76 | 42 | -44 | 26 | 42 | -44 | 26 |
| 1 | 3.35 | 14 | 14 | -20 | 14 | 14 | -20 |
| 1 | 3.38 | -24 | -34 | 60 | -24 | -34 | 60 |
| 1 | 3.74 | 42 | 24 | -22 | 42 | 24 | -22 |
| 1 | 5.27 | 32 | -72 | -38 | 32 | -72 | -38 |

**Table S6.** Fingers, Males > Females

| | | Max Intensity | | | Center of Gravity | | |
|---|---|---|---|---|---|---|---|
| **Voxels** | **Z-Max** | **X** | **Y** | **Z** | **X** | **Y** | **Z** |
| 2658 | 4.66 | 4 | -48 | 42 | 2.86 | -65.3 | 29.2 |
| 1294 | 4.7 | -14 | -24 | 52 | 2.45 | -27.2 | 61.8 |
| 390 | 4.55 | 46 | -52 | 28 | 47.9 | -60.2 | 24.7 |
| 229 | 4.88 | 60 | 2 | 10 | 53.5 | -1.78 | 20.5 |
| 111 | 4.06 | 46 | -40 | 16 | 47.3 | -31.4 | 17.9 |
| 55 | 3.96 | -46 | -54 | 30 | -46 | -56.9 | 27.1 |
| 41 | 4.11 | -38 | -74 | 20 | -40.6 | -69.4 | 18.8 |
| 37 | 4.82 | 36 | -22 | 6 | 36.2 | -21.6 | 3.93 |
| 35 | 4.52 | 12 | -12 | 38 | 11 | -14.3 | 38.3 |
| 33 | 3.7 | 4 | -18 | 40 | 1.64 | -19.2 | 36.1 |
| 29 | 3.43 | 32 | -10 | 46 | 33 | -8.2 | 45.5 |
| 19 | 4.69 | 64 | -2 | 28 | 63.2 | -2.09 | 29.5 |
| 18 | 4.06 | 58 | -32 | 0 | 60.3 | -30.7 | 0.834 |
| 15 | 3.39 | 34 | -74 | 32 | 34.4 | -75.5 | 35.5 |
| 15 | 4.46 | -34 | -22 | 16 | -36.5 | -22.4 | 14.1 |
| 13 | 3.94 | 44 | -50 | 62 | 42.3 | -51.6 | 62.9 |
| 13 | 3.63 | 32 | -2 | 14 | 33.7 | -1.09 | 11.1 |
| 11 | 2.93 | -26 | -82 | 44 | -24.1 | -81.4 | 42.9 |
| 7 | 3.02 | 2 | 6 | 40 | 1.72 | 6.84 | 37.7 |
| 7 | 4.1 | 38 | -34 | 44 | 37.4 | -35.2 | 44.6 |
| 7 | 3.38 | -8 | -36 | 50 | -8.22 | -34.9 | 49.8 |
| 6 | 3.66 | -52 | -66 | 34 | -51.7 | -65 | 32.4 |
| 6 | 3.41 | 0 | -12 | 36 | -1.01 | -8.39 | 34 |
| 5 | 3.91 | 64 | -14 | 12 | 60.1 | -14.4 | 10.4 |
| 4 | 3.73 | 30 | -12 | 8 | 31 | -11.5 | 9.48 |
| 3 | 3.43 | 42 | -2 | 16 | 40.7 | -2.61 | 16 |

Supplementary Tables

| | | | | | | | |
|---|---|---|---|---|---|---|---|
| 3 | 3.3 | 52 | -38 | 14 | 52.6 | -38.6 | 13.4 |
| 2 | 3.59 | 40 | -68 | 44 | 40 | -68.9 | 44 |
| 2 | 3.4 | 22 | -56 | 62 | 23 | -56 | 62 |
| 2 | 3.65 | 54 | -28 | 54 | 53 | -28 | 54 |
| 2 | 3.46 | 40 | -2 | 60 | 40 | -2.99 | 60 |
| 2 | 3.86 | -38 | -22 | -4 | -39 | -22 | -4 |
| 2 | 3.29 | -10 | 0 | 34 | -10 | -0.962 | 34 |
| 2 | 2.6 | 6 | -58 | 36 | 6 | -57 | 36 |
| 1 | 2.85 | -2 | -10 | 30 | -2 | -10 | 30 |
| 1 | 3.71 | -40 | -56 | 22 | -40 | -56 | 22 |
| 1 | 2.43 | 38 | -16 | 38 | 38 | -16 | 38 |
| 1 | 4.4 | 36 | -8 | 20 | 36 | -8 | 20 |
| 1 | 2.45 | -18 | -56 | 18 | -18 | -56 | 18 |
| 1 | 3.32 | 18 | -12 | 40 | 18 | -12 | 40 |
| 1 | 3.25 | 44 | -6 | 14 | 44 | -6 | 14 |
| 1 | 3.32 | 68 | -16 | 14 | 68 | -16 | 14 |
| 1 | 2.83 | 58 | -40 | 14 | 58 | -40 | 14 |
| 1 | 3.74 | 52 | -18 | 12 | 52 | -18 | 12 |
| 1 | 4.36 | 52 | -26 | 50 | 52 | -26 | 50 |
| 1 | 4.08 | -32 | 2 | 6 | -32 | 2 | 6 |
| 1 | 3.64 | 50 | 4 | 54 | 50 | 4 | 54 |
| 1 | 3.83 | 70 | -20 | 4 | 70 | -20 | 4 |
| 1 | 3.75 | 58 | -34 | -4 | 58 | -34 | -4 |
| 1 | 4.2 | 40 | -16 | -8 | 40 | -16 | -8 |

**Table S7.** Ankles, Males > Females

| | | Max Intensity | | | Center of Gravity | | |
|---|---|---|---|---|---|---|---|
| **Voxels** | **Z-Max** | **X** | **Y** | **Z** | **X** | **Y** | **Z** |
| 157 | 5.27 | 58 | 2 | 10 | 60.1 | 2.83 | 17.9 |
| 137 | 4.45 | 42 | -16 | 62 | 39.5 | -17.6 | 50.6 |
| 100 | 4.17 | -16 | -80 | 44 | -13.8 | -78.9 | 35.3 |
| 80 | 4.39 | 30 | -22 | 58 | 27.3 | -24 | 55.4 |
| 17 | 3.5 | 38 | -2 | 62 | 41.2 | -0.763 | 58.7 |
| 14 | 3.81 | 16 | -22 | 68 | 17.8 | -23 | 65.8 |
| 10 | 3.86 | 16 | -80 | 36 | 16 | -79 | 36 |
| 8 | 3.86 | 30 | -10 | 42 | 30.7 | -8.79 | 43.7 |
| 8 | 4.51 | 38 | 6 | 60 | 35.9 | 5.06 | 58.5 |
| 7 | 3.61 | 18 | -70 | 32 | 18.3 | -70.9 | 31.4 |
| 6 | 3.89 | 52 | -2 | 24 | 51.3 | -2 | 24.3 |
| 6 | 3.49 | 60 | -8 | 32 | 57.4 | -7.05 | 30.7 |
| 2 | 4.36 | 48 | 16 | 22 | 48.9 | 16 | 22 |
| 2 | 3.82 | -10 | -64 | 32 | -10 | -64 | 31 |
| 2 | 3.81 | 16 | -24 | 56 | 16 | -25 | 56 |
| 1 | 3.07 | 62 | 12 | 0 | 62 | 12 | 0 |
| 1 | 3.06 | 66 | 2 | 12 | 66 | 2 | 12 |
| 1 | 3.84 | 22 | -70 | 26 | 22 | -70 | 26 |
| 1 | 3.91 | -6 | -84 | 34 | -6 | -84 | 34 |
| 1 | 3.63 | 18 | -66 | 36 | 18 | -66 | 36 |
| 1 | 3.17 | 50 | -10 | 36 | 50 | -10 | 36 |
| 1 | 3.82 | -4 | -82 | 38 | -4 | -82 | 38 |
| 1 | 4.19 | 28 | -70 | 40 | 28 | -70 | 40 |
| 1 | 4.53 | 20 | -80 | 46 | 20 | -80 | 46 |
| 1 | 2.77 | 30 | -20 | 46 | 30 | -20 | 46 |
| 1 | 4.78 | -16 | -60 | 70 | -16 | -60 | 70 |

**Table S8.** Right leg, Females > Males

| | | Max Intensity | | | Center of Gravity | | |
|---|---|---|---|---|---|---|---|
| **Voxels** | **Z-Max** | **X** | **Y** | **Z** | **X** | **Y** | **Z** |
| 494 | 4.56 | -54 | -22 | -22 | -52.1 | -19.9 | -16.2 |
| 95 | 4.19 | -40 | 0 | -34 | -37.9 | -1.44 | -30.1 |
| 21 | 3.38 | -42 | -10 | -30 | -45.4 | -10.3 | -28.6 |
| 8 | 3.78 | -60 | -6 | -4 | -59 | -8.39 | -4.25 |
| 4 | 4.55 | 44 | -22 | -14 | 42.6 | -24.9 | -12.6 |
| 2 | 3.22 | -58 | -8 | -10 | -58 | -8.99 | -10 |
| 1 | 3.03 | -46 | -6 | -36 | -46 | -6 | -36 |
| 1 | 4.09 | 40 | -20 | -14 | 40 | -20 | -14 |
| 1 | 2.9 | -52 | -18 | -10 | -52 | -18 | -10 |
| 1 | 3.34 | -62 | -8 | -10 | -62 | -8 | -10 |
| 1 | 4.7 | 46 | -48 | 4 | 46 | -48 | 4 |
